# Cell-type-specific dysregulated gene expression in the frontal cortex of an Angelman syndrome pig model

**DOI:** 10.1101/2025.08.20.671338

**Authors:** Ashley Coffell, Livia Schuller, Sarah Christian, Scott V. Dindot

**Affiliations:** Department of Veterinary Pathobiology, College of Veterinary Medicine & Biomedical Sciences, Texas A&M University, College Station, TX, USA; Interdisciplinary Graduate Program in Genetics and Genomics, Texas A&M University, College Station, TX, USA; Ultragenyx Pharmaceutical Inc., Novato, CA, USA

**Keywords:** Angelman syndrome, single cell RNA-sequencing, Sus scrofa, pig model, neurodevelopmental disorder

## Abstract

Angelman syndrome is a neurodevelopmental disorder caused by the loss of the maternal allele of the ubiquitin-protein ligase E3A (*UBE3A*) gene. *UBE3A* is imprinted with maternal-allelic expression in neurons of the central nervous system (CNS) and biallelic expression in other cell types. Consequently, in Angelman syndrome, UBE3A is substantially reduced in CNS neurons and reduced by half in other cells. It is unclear how cell-type-specific gene expression in the brain is dysregulated in Angelman syndrome, as previous studies have lacked cell type resolution. Using single nuclei RNA-sequencing, we show that gene expression is dysregulated in neuronal subtypes in the frontal cortex of neonatal pigs with a *UBE3A* maternal deletion. A total of 3,812 unique genes were dysregulated across ten cell type clusters, with most of the dysregulated genes (3,154 genes) in excitatory neurons. Pathway analysis revealed alterations in oxidative phosphorylation, proteasome function, and synaptic function. Additionally, somatostatin (*SST*) — a secreted neuropeptide involved in GABAergic inhibition and synaptic plasticity— was reduced in inhibitory neurons in the cortex and hypothalamus, which correlated with lower circulating SST protein in neonates but not adolescent pigs. Overall, these findings provide greater clarity on the cell-type-specific dysregulation of gene expression and cellular pathways caused by the loss of UBE3A in CNS neurons, expanding our understanding of the molecular pathology in Angelman syndrome.

## INTRODUCTION

Angelman syndrome (OMIM # 105830) is a severe neurogenetic disorder characterized by profound developmental delay, cognitive impairment, motor coordination deficits, absent or reduced speech, ataxic gait, and a uniquely cheerful disposition [1]. It is caused by mutations or epimutations leading to the loss of function or expression of the maternally inherited allele of the ubiquitin-protein ligase E3A (*UBE3A*) gene [2].

The *UBE3A* gene is imprinted in neurons of the central nervous system (CNS), where the maternal allele is expressed and the paternal allele is repressed. In all other cell types, *UBE3A* is biallelically expressed. Transcription of the *UBE3A* paternal allele is repressed by the *UBE3A* antisense (*UBE3A-AS*) transcript, which represents the distal end of the small nucleolar host gene 14 (*SNHG14*) polycistronic transcript [2, 3]. Transcription of *UBE3A-AS* is thought to inhibit transcriptional elongation of the paternal *UBE3A* allele [3]. As such, individuals with Angelman syndrome have substantially reduced *UBE3A* in CNS neurons and approximately fifty percent reduction of *UBE3A* in all other cell types [2, 4].

The UBE3A protein functions as an E3 ligase that polyubiquitinates proteins for degradation by the 26S proteasome [5]. Additionally, UBE3A functions as a transcriptional coactivator of nuclear steroid hormones and can mono-ubiquitinate at least one protein in a non-degradative context [6, 7]. UBE3A has a broad array of protein and gene targets involved in regulating calcium signaling, cytoskeletal dynamics, proliferation, neuronal homeostasis, mitochondrial function, and retinoic acid signaling, amongst other pathways [8-13]. Changes to these pathways following maternal *UBE3A* loss result in deficits in long-term potentiation and synaptic structure and function [14, 15], although the dysregulated genes and molecular pathways underlying the symptoms observed in individuals with Angelman syndrome are poorly understood.

Several studies have examined dysregulated gene expression in Angelman syndrome models using bulk RNA-sequencing or proteomics, which lack cell type resolution [8-10, 12, 16-24]. It is thus unclear how much of the dysregulation stems from neuronal vs non-neuronal *UBE3A* loss in the brain. Our laboratory recently developed a pig model of Angelman syndrome that mirrors the developmental trajectory of the syndrome [25]. Using single nuclei RNA-sequencing, we examined the cell-type-specific dysregulation of genes and pathways in the frontal cortex of neonatal pigs (10-day-old) with a deletion of the maternal *UBE3A* allele. Our results indicate that loss of maternal *UBE3A* dysregulates gene expression primarily in CNS neurons.

## RESULTS

### Experimental design and classification of cell types

Since neurons are predicted to be more severely impacted than other central nervous system (CNS) cell types by the loss of the maternal *UBE3A* allele, we performed single nuclei RNA-sequencing on tissue punches isolated from the frontal cortex of 10-day-old wildtype (n = 4 [male, n = 2; female n =2]) and maternal *UBE3A* deletion (*UBE3A*^-/+^) pigs (n = 4 [male, n = 2; female n =2]) to identify cell-type-specific differentially expressed genes (DEGs). We then performed multiple pathway analyses to identify and characterize dysregulated pathways in each cell type (**Figure 1A, Supplementary File 1**).

**Figure 1.**
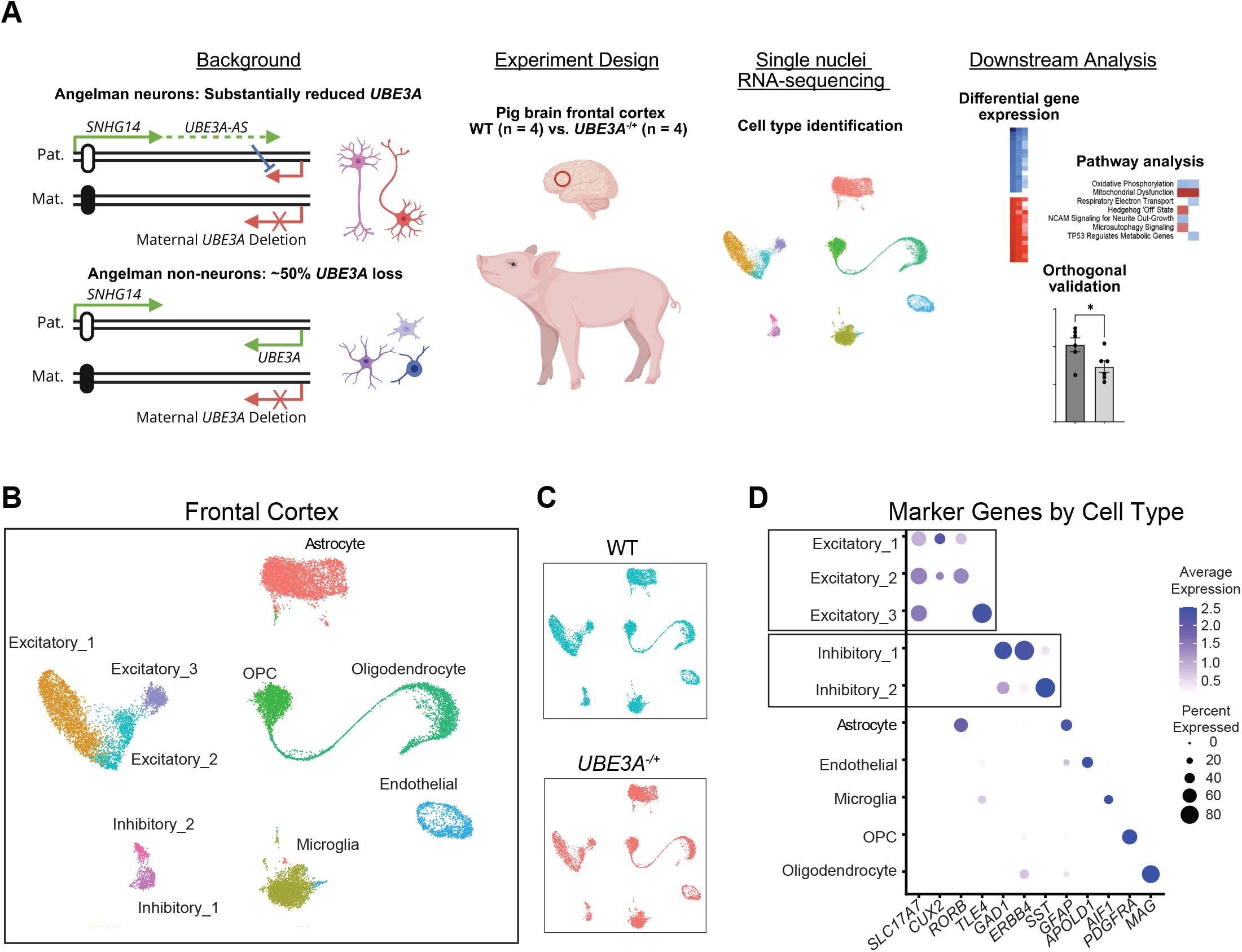
Experimental design and classification of cell types. A. Graphical abstract of the experimental design of the study. B. UMAP of combined single nuclei RNA sequencing data (n = 8), including annotated neuronal and non-neuronal cell types in the frontal cortex. C. UMAP of single nuclei RNA sequencing data for the WT (n = 4) and *UBE3A*^*-/+*^ (n = 4) pigs. D. Percent and average expression of marker genes by cell type and neuronal subtype. Abbreviations: WT, Wildtype; *UBE3A*^*-/+*^, maternal *UBE3A* deletion.

After single nuclei isolation and sequencing, the sequence reads were mapped to the pig NCBI reference genome assembly (*Sus scrofa* 11.1, RefSeq GCF_000003025.6) with custom gene annotations of the *UBE3A* and *UBE3A-AS* transcripts (**see Methods and Supplementary Figure 1**). Analysis of 8,932 WT and 7,942 *UBE3A*^-/+^ cells revealed ten cell types (**Figure 1B**), which clustered independently of genotype (**Figure 1C**), permitting the downstream analysis of differentially expressed genes. The cell types were identified by marker genes expressed in specific cell types, including excitatory neurons (*SLC17A7*), inhibitory neurons (*GAD1*), astrocytes (*GFAP*), oligodendrocytes (*MAG*), oligodendrocyte precursor cells ([OPC] *PDGFRA*), microglia (*AIF1*), and endothelial cells (*APOLD1*) (**Figure 1D, Supplementary Figure 2**) [26-28]. The excitatory and inhibitory neurons were further clustered into different subtypes based on the expression of additional cell-type-specific marker genes (Excitatory_1: *CUX2*, cortical layer 2/3; Excitatory_2: *RORB*, cortical layer 4; Excitatory_3: *TLE4*, cortical layer 5/6; Inhibitory_1: *ERB4*, and Inhibitory_2: *SST*) (**Figure 1D, Supplementary Figure 2**) [29, 30].

**Figure 2.**
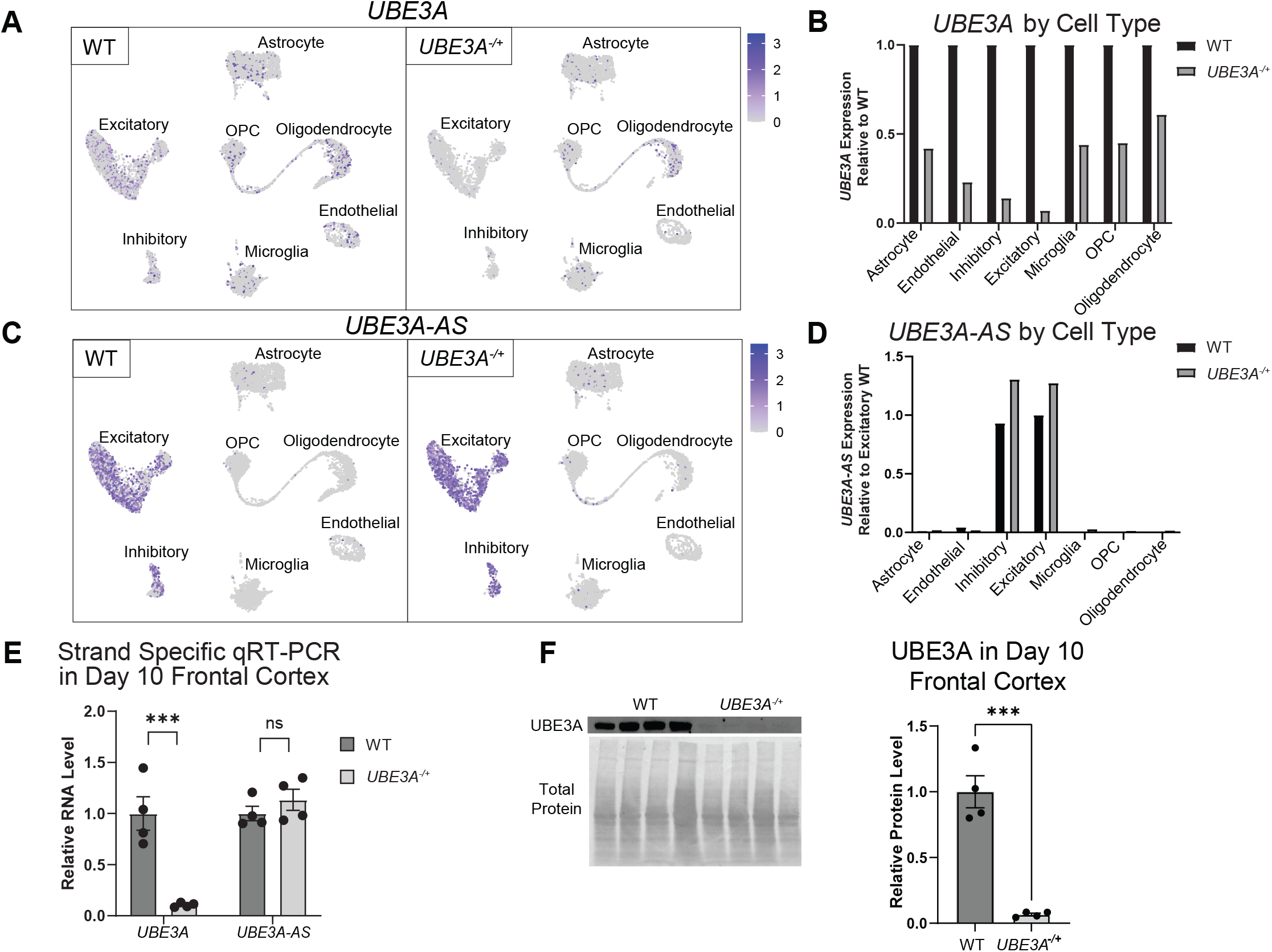
The pig *UBE3A* gene is specifically imprinted in cortical neurons. A. UMAP heatmap of cell type specific *UBE3A* expression in the WT and *UBE3A*^*-/+*^pigs. B. Relative *UBE3A* expression by cell type between WT and *UBE3A*^*-/+*^pigs. Data are presented as the mean *UBE3A* expression (reads per cell/cell type) normalized to WT in each cell type. C. UMAP heatmap of cell type specific *UBE3A-AS* expression in the WT and *UBE3A*^*-/+*^pigs. *D. UBE3A-AS* expression by cell type between WT and *UBE3A*^*-/+*^pigs. Data are presented as the mean *UBE3A-AS* expression (reads per cell/cell type) normalized to WT in excitatory neurons. E. Strand-specific quantitative RT-PCR of *UBE3A* and *UBE3A-AS* expression in the frontal cortex of 10-day-old WT (n = 4) and *UBE3A*^*-/+*^ (n = 4) pigs, normalized to WT. Data are presented as mean ± the standard error of the mean (SEM); Student’s t-test, \*\*\**P* < 0.001; ns, not significant. F. Western blot quantification of UBE3A protein in the frontal cortex of 10-day-old WT (n = 4) and *UBE3A*^*-/+*^ (n = 4) pigs, normalized to total protein. Data are presented as mean ± the SEM; Student’s t-test, \*\*\**P* < 0.001. Abbreviations: UMAP, Uniform Manifold Approximation and Projection; WT, Wildtype; *UBE3A*^*-/+*^, maternal *UBE3A* deletion.

### The pig *UBE3A* gene is specifically imprinted in cortical neurons

Analysis of *UBE3A* and *UBE3A-AS* expression by cell type revealed that *UBE3A* was reduced by 86 – 93% in the neuronal subtypes and by 39 – 77% in the non-neuronal cell types in the *UBE3A*^-/+^ pigs **(Figure 2A and B, Supplementary Figure 1C**). As expected, *UBE3A-AS* expression was limited to the neuronal cell types (**Figure 2C and D**). Interestingly, *UBE3A-AS* was slightly increased in the *UBE3A*^-/+^ pigs, which aligns with a previous finding [31]. We confirmed the imprinting of *UBE3A* in the frontal cortex of 10-day-old WT and *UBE3A*^-/+^ pigs using quantitative RT-PCR and western blot (**Figure 2E and F, Supplementary File 1**).

Altogether, these findings confirm the neuron-specific imprinting of the *UBE3A* gene and expression of the *UBE3A-AS* transcript in pigs.

### Excitatory neurons have the most dysregulated genes in the *UBE3A*^-/+^ pigs

Next, we analyzed each cell type cluster for differentially expressed genes (DEGs). The relative ratios of each cell type were similar and not significantly different between the WT and *UBE3A*^-/+^ pigs (**Figure 3A**), indicating that loss of maternal *UBE3A* does not affect the differentiation of cells in the frontal cortex. Differential expression analysis identified 3,992 DEGs, including 1,527 downregulated and 2,465 upregulated genes in the *UBE3A*^-/+^ pigs with a Bonferroni-adjusted P-value < 0.05 (**Figure 3B, Supplementary File 2**). One hundred and eighty genes showed dual patterns of dysregulation across the cell types (i.e., downregulated and upregulated), reducing the number of unique DEGs to 3,812 (**Supplementary Figure 3, Supplementary File 3**). Excitatory neuron clusters 1 and 2 had the largest number of DEGs, followed by astrocytes. Overall, most DEGs (3,154/3,812 = 82.7%) were detected in the excitatory neuronal clusters (**Supplementary File 3)**. The top upregulated and downregulated genes, sorted by fold-change and adjusted *P*-value, are shown in **Figure 3C and Supplementary File 4**. As expected, *UBE3A* was one of the top downregulated genes in the neuronal clusters, except for the Inhibitory_2 cluster (**Figure 3C**). Using quantitative RT-PCR and bulk RNA-sequencing, we confirmed the dysregulation of several genes in the *UBE3A*^-/+^ cortex in two age groups (10 and 140-day-old [**Supplementary Figure 4, Supplementary File 5**]), including the downregulation of the integral membrane protein 2B (*ITM2B)* and somatostatin (*SST*) genes. Since larger clusters will often yield a greater number of DEGs, we also performed a perturbation analysis, which regresses the cluster size to normalize the number of DEGs per cluster [32]. The results confirmed that the excitatory and inhibitory neurons are the most impacted cell types in the *UBE3A*^-/+^ pigs (**Figure 3D**). These findings show that almost all the dysregulated genes are in neuronal cells in the *UBE3A*^-/+^ pigs, with excitatory neurons having the largest number of DEGs.

**Figure 3.**
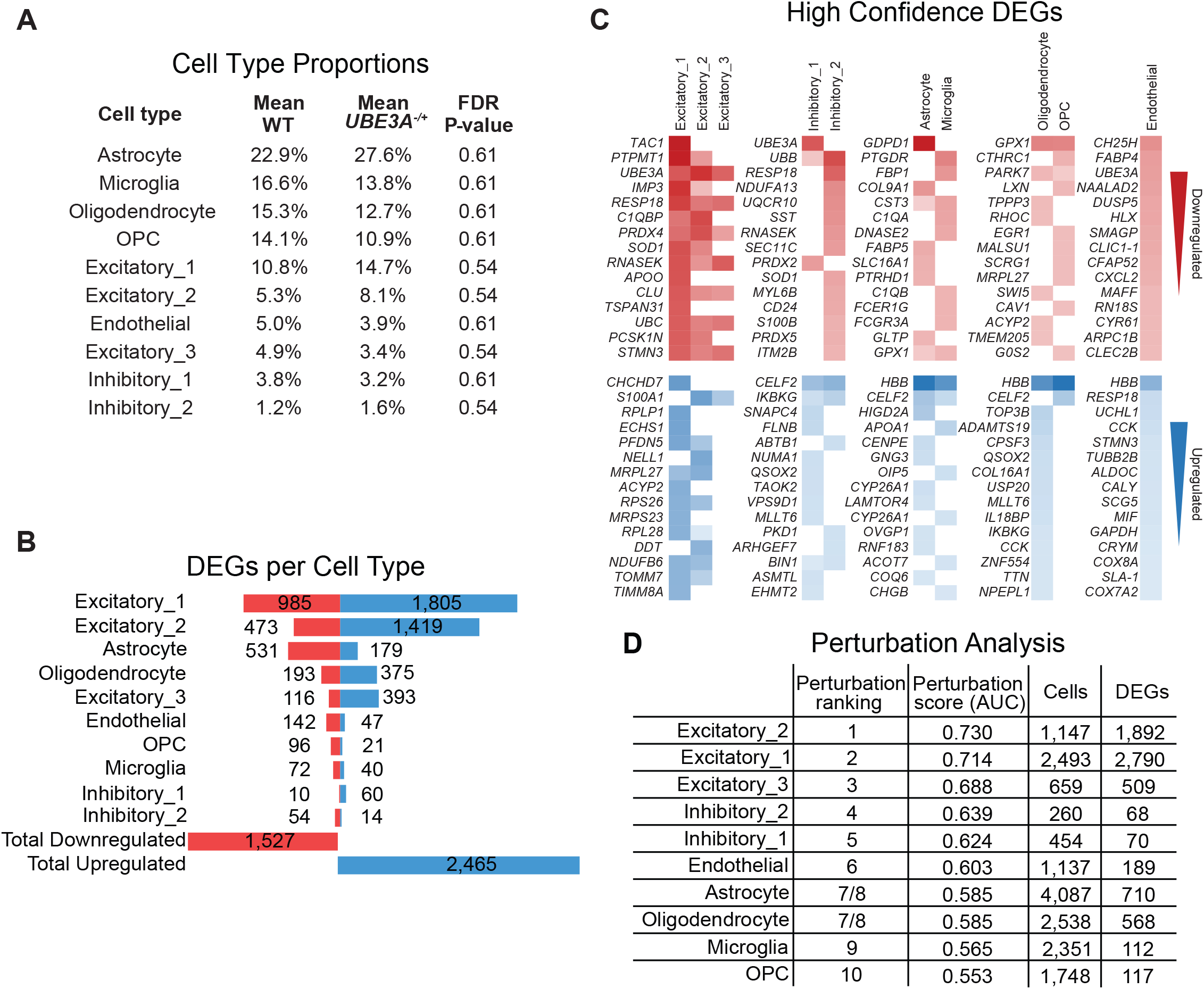
Excitatory neurons have the most dysregulated genes in the *UBE3A*^-/+^ pigs. A. Proportions of cell types in the WT and *UBE3A*^-/+^ pigs. B. DEGs identified in the *UBE3A*^-/+^ pigs by cell type. C. Top 15 upregulated and downregulated high confidence DEGs per excitatory neurons, inhibitory neurons, glia, oligodendrocytes and precursor cells, and endothelial cells. Color corresponds to P-value; blue = upregulated; red = downregulated. *D*. Perturbation analysis of DEGs by cell type. Abbreviations: DEGs, differentially expressed genes; WT, Wildtype; *UBE3A*^*-/+*^, maternal *UBE3A* deletion.

**Figure 4.**
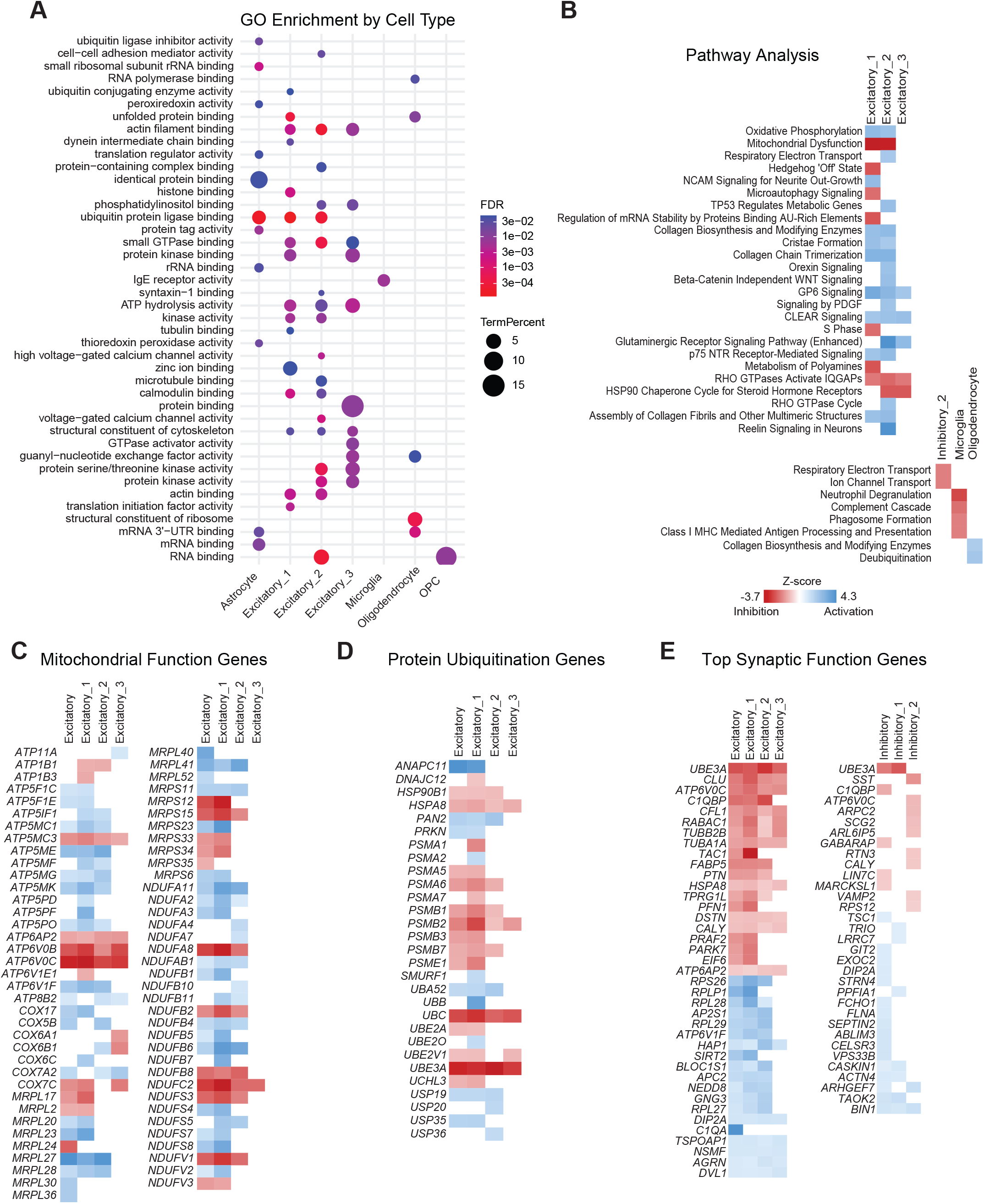
Multiple pathways are dysregulated in excitatory neurons in the *UBE3A*^-/+^ pigs. A. Molecular function gene ontology (GO) enrichment terms by cell type. B. Pathway analysis results are shown by cell type for pathways with significant -log(P-values) (> 1.3) and significant Z-scores (< -2 or > 2). Red denotes pathway inhibition; blue denotes pathway activation. C. Mitochondrial function DEGs with log2 fold change > 1 for the combined excitatory neuron analysis, Excitatory_1, Excitatory_2, and Excitatory_3 are shown. *D*. Protein ubiquitination DEGs with log2 fold change > 1 for the combined excitatory neuron analysis, Excitatory_1, Excitatory_2, and Excitatory_3 are shown. E. The top 20 downregulated and top 20 upregulated DEGs (log2 fold change > 1) for synaptic function are shown for the combined excitatory analysis and the three excitatory subclusters, and all synaptic function genes are shown for the combined inhibitory analysis and the two inhibitory neuron subclusters. Abbreviations: DEGs, differentially expressed genes.

### Multiple pathways are dysregulated in excitatory neurons in the *UBE3A*^**-/+**^ pigs

To identify molecular and cellular pathways dysregulated in each cell type of the *UBE3A*^-/+^ pigs, we first performed a molecular function gene ontology (GO) analysis of the DEGs in each cell type using DAVID (**Figure 4A**). The term “ubiquitin protein ligase binding” had one of the lowest FDR values in the Astrocytes, Excitatory_1, and Excitatory_2 clusters. Significant GO terms were also observed for “unfolded protein binding,” “actin filament binding,” and “small GTPase binding” as well as the modulation of RNAs (e.g., “mRNA 3’−UTR binding,” “mRNA binding,” and “RNA binding”). We next performed a Qiagen Ingenuity Pathway Analysis of high-confidence DEGs (P-adj > 0.05; log2FC > ±1) (**Figure 4B**), which generates a pathway score (activation or inhibition) based on the measured upregulated or downregulated genes in a pathway. Results showed multiple dysregulated pathways for the excitatory neuronal clusters, whereas only a few dysregulated pathways were detected in the Inhibitory_2, Microglia, and Oligodendrocytes clusters, and no dysregulated pathways in the remaining clusters (**Figure 4B, Supplementary File 6**). The top dysregulated pathways in the neuronal clusters involved mitochondrial function (Oxidative Phosphorylation, Mitochondrial Dysfunction, and Respiratory Electron Transport), Hedgehog ‘Off’ State, NCAM Signaling for Neurite Out-Growth, Microautophagy Signaling, and TP53 Regulates Metabolic Genes. Perhaps not surprisingly, Excitatory neuron clusters 1 and 2 contained the largest number of dysregulated pathways and the most significant P-values and Z-scores (**Supplementary File 6**). Further analysis of the high-confidence DEGs revealed an enrichment of dysregulated genes involved in mitochondrial function, protein ubiquitination, and synaptic function (**Figure 4C – E**). The top 40 DEGs for Excitatory Synaptic Function are shown in **Figure 4E**. Altogether, these findings indicate that excitatory neurons are the most highly affected cell type in the frontal cortex of neonatal *UBE3A*^-/+^ pigs, with several dysregulated genes in pathways involving protein ubiquitination, mitochondrial function, and synaptic function.

### Somatostatin (*SST*) expression is reduced in *UBE3A* maternal deletion (*UBE3A*^*-/+*^) pigs

In the single nuclei RNA-sequencing, we found that *SST* was reduced in the Inhibitory_2 cluster and that the somatostatin receptors *SSTR1, SSTR2*, and *SSTR3* were also differentially expressed in excitatory neurons of the frontal cortex of the *UBE3A*^-/+^ pigs (**Supplementary Figure 4, Supplementary File 2**), suggesting that the loss of maternal *UBE3A* expression disrupts this pathway. Somatostatin is a neuromodulator released from a subpopulation of inhibitory GABAergic neurons, where it functions to increase the strength of the inhibitory signal when co-released with GABA [33-35]. Outside of the CNS, it also regulates digestive hormones in the gastrointestinal tract. To confirm these findings and evaluate SST as a potential biomarker for Angelman syndrome, we analyzed *SST* expression in different brain regions and in the serum and cerebrospinal fluid (CSF) of *UBE3A*^-/+^ and WT pigs at different ages.

Quantitative RT-PCR analysis showed that *SST* expression was significantly reduced in the cortex of 10-day-old *UBE3A*^-/+^ pigs relative to WT pigs (26 % reduction, *P* = <0.0001, Mixed-effect linear regression model [**Figure 5A**]), including the frontal, temporal, and occipital regions (*P* = 0.002; *P* = 0.02; *P* = 0.02). SST expression was slightly reduced in the parietal region but not significantly different (*P* = 0.2). Similarly, *SST* expression was slightly reduced in the frontal cortex of embryonic day 103 *UBE3A*^-/+^ pigs (14% reduction, *P* = 0.3, Student’s t-test [**Figure 5B**]) and significantly reduced in the frontal cortex of 60-day-old (27% reduction, *P* = 0.002, Student’s t-test [**Figure 5B**]) and 140-day-old *UBE3A*^-/+^ pigs (26% reduction, *P* = 0.04, Student’s t-test [**Figure 5B**]), indicating that dysregulation of *SST* persists into adult animals. To determine if reduced *SST* expression in the brain correlates with lower levels in circulation, we quantified SST protein levels in 10-day-old pigs using an ELISA assay. Results showed that the average serum SST protein levels were significantly lower in the *UBE3A*^*-/+*^ pigs compared to the WT pigs (WT mean = 190.6 pg/mL; *UBE3A*^*-/+*^ mean = 85.2 pg/mL, *P* = 0.04, Student’s t-test [**Figure 5C**]); however, plasma SST protein levels in 120-day-old pigs were only slightly lower and not significantly different (WT mean = 1,123 pg/mL; *UBE3A*^-/+^ mean = 908.5 pg/mL, *P* = 0.3, Student’s t-test [**Figure 5C**]). SST protein levels in the CSF were similar and not significantly different between the *UBE3A*^*-/+*^ and WT pigs (WT mean = 38.1 pg/mL; *UBE3A*^-/+^ mean = 37.2 pg/mL, *P* = 0.7, Student’s t-test [**Figure 5C**]). Likewise, bulk RNA-sequencing showed that SST expression was not reduced in the spinal cord of *UBE3A*^*-/+*^ pigs compared to the WT pigs (**Supplementary File 7**).

**Figure 5.**
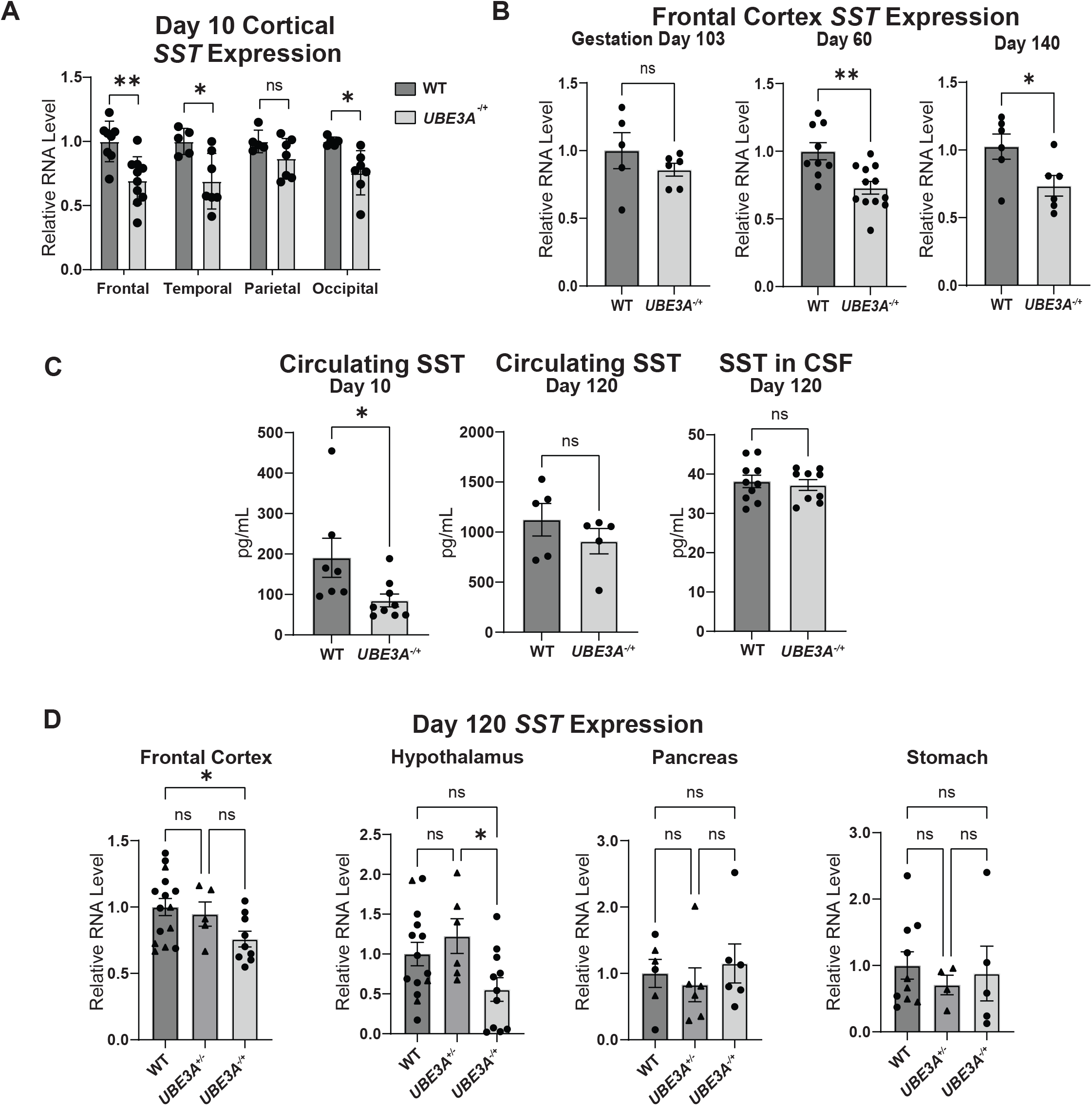
Somatostatin (*SST*) expression is reduced in *UBE3A* maternal deletion (*UBE3A*^*-/+*^) pigs. A. Quantitative RT-PCR of *SST* in day 10 frontal, temporal, and occipital cortex, but not parietal cortex (WT n = 5-8, *UBE3A*^-/+^ n = 7-10). Data are presented as mean ± the standard error of the mean (SEM); Student’s t-test and mixed-effect linear regression model \**P* < 0.05; \*\**P* < 0.01; ns, not significant. B. Quantitative RT-PCR of *SST* in gestation day 103 (WT n = 5, *UBE3A*^-/+^ n = 6) and postnatal day 60 (WT n = 9, *UBE3A*^-/+^ n = 12) and 140 (WT n = 6, *UBE3A*^-/+^ n = 6). Data are presented as mean ± the standard error of the mean (SEM); Student’s t-test, \**P* < 0.05; \*\**P* < 0.01; ns, not significant. C. SST protein measured by ELISA in the serum of 10-day-old (WT n = 7, *UBE3A*^-/+^ n = 9), plasma of 120-day-old (WT n = 5, *UBE3A*^-/+^ n = 5) and CSF measured in 120-day olds pigs (WT n = 10, *UBE3A*^-/+^ n = 9). Data are presented as mean ± the standard error of the mean (SEM); Student’s t-test, \**P* < 0.05; ns, not significant. *D. SST* qRT-PCR results in frontal cortex, hypothalamus, pancreas, and stomach in adolescent (120-day-old) WT (n = 6-15), *UBE3A*^*+/-*^ (n = 4-6), and *UBE3A*^*-/+*^ (n = 5-11) animals. Data are presented as mean ± the standard error of the mean (SEM); analysis of variance (ANOVA) followed by a Tukey post-hoc test, \**P* < 0.05; ns, not significant. Abbreviations: Enzyme-Linked Immunosorbent Assay = ELISA; Wildtype = WT; *UBE3A* paternal deletion = *UBE3A*^*+/-*^; *UBE3A* maternal deletion = *UBE3A*^*-/+*^.

Serum SST is derived from both the hypothalamus and gastrointestinal (GI) tract [36, 37]. To determine if the reduced serum SST protein expression in the *UBE3A*^*-/+*^ pigs is due to the loss of UBE3A in the hypothalamus or haploinsufficiency in the GI tract, we examined SST expression in the frontal cortex, hypothalamus, pancreas, and stomach of 120-day-old WT, *UBE3A*^-/+^, and *UBE3A*^+/-^ pigs. In the frontal cortex and hypothalamus, *SST* expression was similar and not significantly different between the WT and *UBE3A*^+/-^ pigs, but it was reduced in the *UBE3A*^-/+^ pigs, with several pigs having almost no detectable *SST* expression (**Figure 5D**). In contrast, *SST* expression was similar and not significantly different across all three genotypes in the pancreas and stomach (**Figure 5D**). Thus, loss of *UBE3A* expression in the hypothalamus of the *UBE3A*^-/+^— but not the *UBE3A*^+/-^— pigs leads to a reduction in *SST* expression in the hypothalamus, which in turn leads to reduced SST protein in circulation. This effect, however, is highly variable among animals and at different ages, indicating that other factors also influence the expression and, perhaps, secretion of SST.

## DISCUSSION

Using single nuclei RNA-sequencing, we identified 3,812 differentially expressed genes (DEGs) in the frontal cortex of the neonatal *UBE3A*^-/+^ pigs. Excitatory and inhibitory neurons were the main cell types affected by the loss of maternal *UBE3A*, consistent with the neuron-specific imprinting of *UBE3A*. Pathway analyses revealed that these DEGs were enriched in several pathways, including the production of ATP in mitochondria, protein ubiquitination, and synaptic function.

Our findings show that the majority of DEGs (83%) and affected pathways were detected in excitatory neurons. Likewise, the perturbation analysis, which corrects for the abundance of cells in the analysis, indicated that both the excitatory and inhibitory neurons were the most affected cell types. This large effect on gene dysregulation in neurons — and not non-neuronal cells — is consistent with the neuron-specific imprinting of *UBE3A*, further supporting the notion that loss of *UBE3A* expression in CNS neurons is the primary cause of the symptoms associated with Angelman syndrome. Indeed, DEGs were detected in non-neuronal cells, but our pathway analyses only detected a few perturbed pathways, suggesting that UBE3A is haplosufficient in these cell types. This conclusion is supported by the lack of overt phenotypes in both animal models and people with a paternally inherited deletion of *UBE3A*.

Our findings indicate that loss of maternal *UBE3A* leads to the dysregulation of several genes involved in ATP production, protein ubiquitination, and synaptic function in neurons is consistent with the known roles of UBE3A. Moreover, our findings correlate with the findings of Pandya et al. [38], who used a proteomics approach to identify dysregulated proteins and pathways in mouse, rat, and human models of Angelman syndrome. The concurrent findings of the transcriptomic and proteomic data suggest these are the focal pathways disrupted with maternal *UBE3A* loss. It remains unclear if the changes in gene expression precede protein abundance alterations or are secondary responses to protein alterations. Unlike Pandya et al., we did not observe dysregulated gene expression of aminoacyl tRNA synthetases (*AIMP1, MARS1, YARS, WARS*), suggesting that changes in these protein abundances occur post-transcriptionally (**Supplementary File 2**).

Our results showing that *SST* expression is reduced in the cortex and hypothalamus of the *UBE3A*^-/+^ pigs are also consistent with prior studies. SST serves as a neurotransmitter in GABAergic neurons, where it is released concurrently with GABA, strengthening the inhibition signal [39]. Our finding that *SST* is reduced in *SST*+ inhibitory neurons is consistent with the prior observations that (1) inhibitory signal deficits are found in a mouse model of Angelman syndrome and (2) the number of SST+ neurons is not affected [40]. Kim et al. [41] previously found *SST* reduced in the hippocampus of *UBE3A-null* mice. SST is crucial for long-term potentiation, plasticity, and motor learning and may contribute to the cognitive deficits observed in individuals with Angelman syndrome [42-44]. Further, loss of SST in the dentate gyrus has been associated with seizure activity, and SST has been investigated for antiepileptic properties [45, 46]. Cortical SST has been shown to be independent of sex hormones, but hypothalamic SST is regulated at the RNA level through transcriptional impacts of sex hormones [47, 48]. In our data, we did not observe an effect due to sex; however, our sample size was limited and the animals investigated were prepubescent (**Supplementary File 9**).

We acknowledge several limitations of the study. First, although we confirmed several DEGs in older animals, the results of this study are mainly limited to the frontal cortex of neonatal animals, and thus, additional studies are needed to evaluate the dysregulation of genes in additional brain regions and at different ages. Second, we had limited success in validating several DEGs using the bulk tissue methods (i.e., quantitative RT-PCRs and bulk RNA-sequencing). Whether this is due to contamination by other cell types or false positives associated with the analysis warrants further investigation. Third, the single-nuclei isolation protocol and distribution of cells within the tissue inevitably lead to variability among the cell type proportions, with cells present at higher proportions having greater statistical power to distinguish differential gene expression. For instance, inhibitory neurons, which were the least abundant cell type in this study, also had the fewest number of DEGs. In the perturbation analysis, however, inhibitory neurons had the highest perturbation score after excitatory neurons, indicating that a high proportion of genes are dysregulated in this cell type but likely missed due to their abundance. We attempted to overcome this situation by combining the excitatory and inhibitory clusters (**Supplementary File 10**). Indeed, additional DEGs were identified by combining the cell types, but DEGs were also lost (e.g., *SST* in Inhibitory_2), highlighting the utility of the cell type clustering approach to identify DEGs.

In conclusion, we show that loss of maternal *UBE3A* expression mainly dysregulates gene expression in neonatal cortical neurons of a pig model of Angelman syndrome. Our study and findings highlight the use of a single cell approach to identify dysregulated genes and pathways, providing greater clarity on the cell-type-specific dysregulation caused by the loss of maternal *UBE3A* expression, which is subject to a complex cell-type-specific pattern of regulation.

## METHODS

### Study animals

The study animals included wild-type pigs (*UBE3A*^+/+^), pigs with maternally inherited 97 kilobase deletion of the *UBE3A* gene (*UBE3A*^-/+^ [NC_010443.5:g.141887260_141984697del]), and pigs with a paternally inherited 97 kilobase deletion of the *UBE3A* gene (*UBE3A*^+/-^) [25]. The study animals were housed in small groups (approximately 3-4 pigs) in climate-controlled rooms under twelve-hour light cycles. Genotyping was performed through a polymerase chain reaction (PCR), as previously described [25]. Feed was provided ad libitum through day 40 and afterward, was fed at 2–4% daily of body weight. Animals past weaning age (approximately 21 days) were food deprived the night before necropsying. Animals did not receive treatment for Angelman syndrome phenotypes. All animal work was conducted under the guidance, supervision, and approval of the Texas A&M University Institutional Animal Care and Use Committee. A table of all animals used in this study is available in **Supplementary File 1**.

Postnatal animals were sedated (intramuscular tiletamine) and anesthetized (isoflurane ventilation), then euthanized via cardiac perfusion with phosphate-buffered saline. Animals used in the bulk RNA-sequencing were euthanized with phenytoin/pentobarbital. Fetal pigs were collected by a caesarean-section while the sow was sedated (intramuscular delivery of ketamine, butorphanol, and midazolam) and anesthetized (isoflurane ventilation). Areflexic fetal pigs were euthanized with 5 mL of intravenous potassium chloride prior to tissue collection. Brain region-specific and body tissue samples were taken in 4 mm punches, flash-frozen in liquid nitrogen, and then stored at -80°C until use.

### Single nuclei library preparation

Brain region-specific samples were taken in 4 mm punches, flash-frozen in liquid nitrogen, and then stored at -80°C until use. Using the 10X Genomics Nuclei Isolation Kit (#PN-1000493, 10X Genomics, CA, USA), nuclei were isolated from 25 mg frontal lobe cortex tissue from four WT and four *UBE3A*^*-/+*^ 10-day-old piglets. Each genotype was comprised of two males and two females. The 10X Chromium Connect was used for automated Single Cell Gene Expression 3’ v3 library preparation (#PN-1000075, 10X Genomics), followed by sequencing on Illumina NovaSeq 6000, XP Workflow (Illumina, CA, USA).

### Single nuclei RNA-sequencing computational analysis

Cellranger (7.2.0) mkref was used to create a reference genome from NCBI *Sus scrofa 11*.*1* (GCF_000003025.6) plus a custom annotation of *UBE3A* and *UBE3A-AS* (**Supplementary Figure 1**). Cellranger count was used for generating count matrices with default settings applied, except for introns exclusion with –include-introns=false. SoupX (1.6.2) ambient RNA correction was applied to Cellranger count outputs. Scater (1.0.3) was used to calculate median absolute deviations (MADs) for thresholding feature counts per sample. Using Seurat (5.1.0), barcodes were excluded that had feature counts >5 or <3 MADs per sample and ≥5% mitochondrial genes (annotated with KEF22-prefix in pigs). After initial Seurat CCA integration (dims = 30, resolution = 0.5), clusters with an average of <900 features per cell were removed. After filtering and computational analysis, a final count of 16,874 cells (8,932 WT; 7,942 *UBE3A*^*-/+*^) was retrieved. Final figures and analysis were then generated with Seurat CCA integration (dims = 20, resolution = 0.5). scType was used to generate initial cell type labels for each cluster and labels were confirmed to agree with published markers [49]. Based on the clustering results, the neuronal clusters were further sub-typed into *ERBB2*+ (Inhibitory_1) and *SST*+ (Inhibitory_2) inhibitory neurons, and layer 2/3 (*CUX2*+; Excitatory_1), layer 4 (*RORB*+; Excitatory_2), and layer 5/6 (*TLE4*+; Excitatory_3) excitatory neurons consistent with published markers [26-28]. Differential expression analysis was performed using the FindMarkers function in Seurat using the default settings (Wilcox test with a threshold of log2FC of 0.1) and significance was adjusted for multiple comparisons using a Bonferroni-adjusted P-value of <0.05. Further analyses were performed by applying Augur (1.0.3) perturbation analysis and Speckle (1.6.0) cell type proportion analysis. Genes reported with *LOC* prefix have not been assigned a gene symbol by NCBI. Average *UBE3A* and *UBE3A*-*AS* expression values were values returned with the Seurat AverageExpression function. For illustrating *UBE3A* expression, values were displayed relative to each wildtype cell type to compare relative *UBE3A*^-/+^ vs wildtype expression. For illustrating *UBE3A-AS* expression, values were displayed relative to Excitatory wildtype to compare expression by cell type. For the list of high confidence genes, data was sorted in Qiagen Ingenuity Pathway Analysis, where only genes with expression > 0 and with known human orthologues were kept, then sorted by P-value and average log2FC. A psdeuobulk analysis was also performed in Seruat, where each animal served as a tissue replicate, and differential expression analysis was performed with DESeq2.

### Custom annotation of *UBE3A* and *UBE3A-AS*

The custom annotation of *UBE3A-AS* was generated from frontal cortex bulk RNA-sequencing using methods previously described [50]. Briefly, FASTQ sequences generated from strand-specific poly-A enriched RNA were aligned to Sus scrofa 11.1 with Hisat2 (2.1.0), then aligned SAM sequences were then converted to binary BAM sequences, indexed, and sorted using samtools. Aligned sequences were filtered remove non-uniquely aligned reads. The 5’ and 3’ reads were split using samtools. Transcript assemblies were generated using Stringtie (1.3.4.d), with the setting: -g 10. Single-exon transcripts and transcripts with non-canonical splice-sites were filtered using gffread (GFF utilities). Transcript units were defined as overlapping transcripts expressed from the same strand of a transcriptional unit and sharing a common terminal exon. Cellranger mkref was used to create a custom reference with the *UBE3A*/*UBE3A*-*AS* overlapping exons removed, which correspond to the following coordinates:

chr1:141984591-141987465; chr1:141974996-141975160; chr1:141925289-141925484; chr1:14188718-141887772.

### Quantitative reverse transcription polymerase chain reaction (qRT-PCR)

RNA from flash-frozen samples was isolated with Qiagen RNeasy Plus kit (#74134, Thermo Fisher Scientific, MA, USA). RNA was quantified with Qubit RNA BR Assay kit (Thermo Fisher Scientific, #Q10210) and a Qubit 4 Fluorometer. First-strand cDNA synthesis was performed with the SuperScript IV kit (Thermo Fisher Scientific, #18091200) with the Oligo(dT)20 template for poly-A transcript enrichment. For *SST* in all tissues except the pancreas, TaqMan reactions (TaqMan Gene Expression Master Mix, #4369016, Thermo Fisher Scientific) were performed according to the manufacturer’s instructions and normalized to *RPL4* levels. The TaqMan assays used were *SST* (Ss03391856_m1) and *RPL4* (Ss03374067_g1). For *UBE3A* and *UBE3A-AS* in frontal cortex and *SST* in pancreas, SYBR green qRT-PCR reactions (PowerUp SYBR Green Master Mix for qPCR, #A25742. Thermo Fisher Scientific) were performed according to manufacturer’s instructions and used *PPIA* for measuring relative expression. Primers sequences used are available in **Supplementary File 11**). Reactions were run in triplicate, and replicate values were excluded if the standard deviation exceeded greater than a quarter of a cycle. Twenty μL reactions with 2 μL of template (equivalent to 8 ng or 16 ng of RNA) were run on a Bio-Rad CFX96 Touch Real-Time thermocycler (Bio-Rad, CA, USA). The parameters for the SYBR assays are as follows: 50°C for 2 minutes, 95°C for 2 minutes, then 40 cycles of 95°C for 15 seconds, 60°C for 1 minute, with readings taken at the end of every 60°C step. The parameters for the TaqMan assays are as follows: 50°C for 2 minutes, 95°C for 10 minutes, then 40 cycles for 95°C for 15 seconds, 60°C for 1 minute, with readings taken at the end of every 60°C step. Results were analyzed with Bio-Rad CFX Maestro software.

### Western blot and densitometry

Tissue samples were lysed in NP40 buffer (comprised of 1% Nonidet P40 and 0.01% SDS in 100mM Tris-HCl [pH 7.2]) with Roche protease inhibitor (#11697498001, Sigma-Aldrich, MA, USA). Lysates were mixed with 4x Laemmli buffer (#1610747, Bio-Rad,) with 10% β-mercaptoethanol added and heated at 95°C for 10 minutes. Proteins were separated on Bio-Rad 10% Mini-PROTEAN TGX precast gels (#4561033, Bio-Rad) at 30 V for 30 minutes, then 100 V for 45 minutes, and transferred to a nitrocellulose membrane using the Bio-Rad Trans-Blot Turbo System. The membrane was blocked in a buffer comprised of 5% nonfat dry milk in 0.1M Tris-buffered saline with 0.1% Tween-20 (TBST), rotating for 1 hour at room temperature, then incubated overnight rotating at 4°C with mouse anti-UBE3A antibody (#611416, BD Biosciences, CA, USA) diluted 1:1,000 in the blocking buffer. After washing in TBST for three washes of 15 minutes, the membrane was incubated with goat anti-mouse HRP-conjugated secondary antibody (#626520, Thermo Fisher Scientific) diluted 1:10,000 in the blocking buffer for 1 hour at room temperature, rotating. Protein detection was performed using Bio-Rad Clarity Western ECL Substrate (#1705061, Bio-Rad) and visualized on a Bio-Rad ChemiDoc Imaging System. Densitometry analysis was performed in ImageJ. UBE3A protein levels were normalized to the amount of Ponceau S stain (#K793, VWR, OH, USA) total protein per lane.

### Bulk RNA-sequencing

Samples were flash-frozen in liquid nitrogen and then stored at -80°C until use. RNA was extracted from 20 mg of tissue using the Qiagen RNeasy kit (#74104, Thermo Fisher Scientific) following the manufacturer’s protocol. RNA was eluted from the columns using 60 μL of nuclease-free water. RNA quantity was verified using the Qubit RNA BR assay (#Q10210, Thermo Fisher Scientific). RNA quality was checked using an Agilent TapeStation. (Agilent, CA, USA) RNA libraries were prepared from 1 μg of total RNA with RIN > 6.1 by TIGSS using the TruSeq mRNA stranded kit (#20020595, Illumina). Paired-end, 150 base pair sequencing on the NovaSeq 6000 at a depth of 10-20 million reads per sample was performed. Qiagen IPA RNA Portal was used for Ensembl *Sscrofa11*.*1*.*105* reference alignment and differential expression analysis with default parameters. Gene names with the *ENSSSCG* prefix represent unnamed genes or genes in which a human ortholog has not been identified. Ensemble gene IDs were converted to RefSeq gene IDs using conversion resources from the HGNC Comparison of Orthology Predictions (HCOP) tool (genenames.org/tools/hcop).

### Gene ontology and pathway analysis of dysregulated genes

DAVID functional annotation was performed with default settings for gene ontology. Dysregulated pathways were identified using QIAGEN Ingenuity Pathway Analysis [51]. The DEGs were entered by cell type. Enriched pathways represent pathways where there are a greater number of genes differentially expressed than would be expected by random chance, while pathway activation and inhibition scores are predicted based on the observed up- or downregulation of the genes. **Figure 4B** shows the top pathways with both significant Z-scores and P-values for high confidence DEGs (P-adj > 0.05; log2FC > ±1). Some pathways not relevant to CNS tissue and pathways enriched with genes inherent to variation in single cell data (ribosomal genes) were excluded but are available in **Supplementary File 6**. Pathway analysis for all DEGs (P-adj > 0.05) is included in Supplementary File 8). Genes related to synaptic function were determined with SynGO.

### Enzyme-linked immunosorbent assay (ELISA)

Blood and CSF samples were taken prior to euthanasia. Blood samples from 10-day-old piglets were collected into serum separator tubes (#367814, BD Biosciences), while blood samples from 120-day-old animals were collected into K2 EDTA tubes (#367861, BD Biosciences). Blood samples were centrifuged at 1200 x g for 10 minutes, aliquoted, and stored at -80**°**C. CSF was taken from suboccipital cisterna magna collection. A competitive ELISA from Novus Bio specific for SST (#nbp2-80269, Novus Bio, CO, USA) was used to quantify SST, according to the manufacturer’s instructions. Absorbance at 450 nm was measured with an Agilent BioTek Cytation5. Absorbance values were interpolated to pg per mL by a standard curve in GraphPad Prism with an asymmetric five-parameter logistic equation fit test. The standard curve was run in duplicate, and samples were run in duplicate or triplicate. Triplicate measurements exceeding 10% coefficient of variation were excluded.

### Statistical Analyses

Statistical analyses were performed in GraphPad Prism (GraphPad, CA, USA) or JMP (JMP Statistical Discovery, NC, USA). For comparison between two groups, a Student’s t-test was performed. For comparisons between three groups, an ANOVA (Analysis of Variance) test with a Tukey’s multiple comparison’s post-hoc test was used. For multiple comparisons with different tissues from an individual animal, Mixed-Effect Logistic Regression model was used to account for repeated measures.

## Acknowledgments

The authors would like to thank the assistance of Dr. Andrew Hillhouse and the Texas A&M Institute for Genome Sciences and Society (TIGSS) for assistance with the RNA-sequencing and study design.

