## Supplemental Figures for "Cell-type-specific dysregulated gene expression in the frontal cortex of an Angelman syndrome pig model"

### Supplementary Figures and Legends

#### Supplementary Figure 1

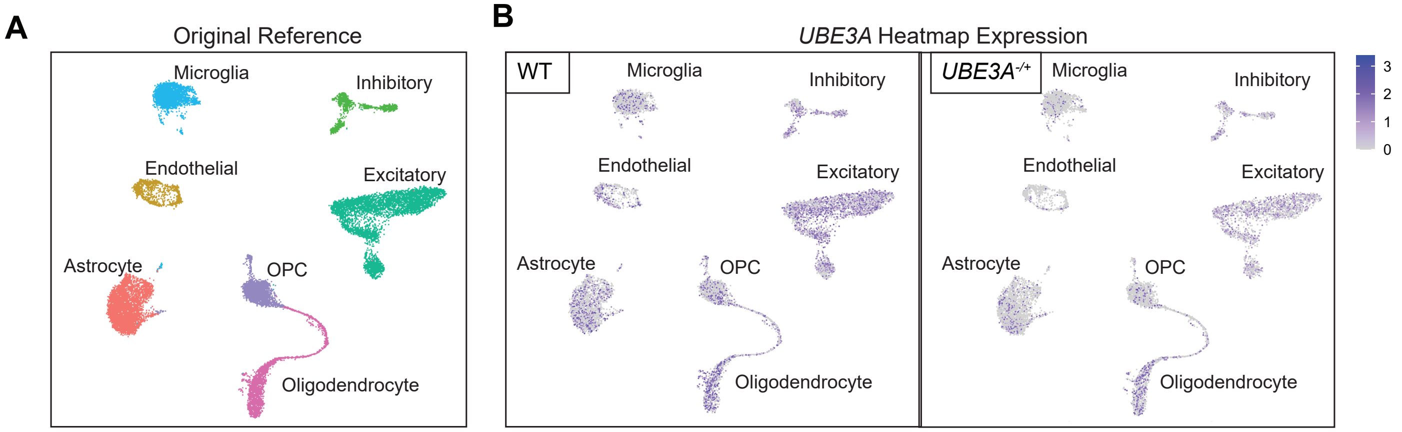


**C**

Relative *UBE3A* Expression
(*UBE3A*-/+/WT)

| | Original Reference | No Overlapping Exons |
|---|---|---|
| Astrocyte | 46% | 42% |
| Endothelial | 27% | 23% |
| Inhibitory | 64% | 14% |
| Excitatory | 50% | 7% |
| Microglia | 46% | 44% |
| OPC | 47% | 45% |
| Oligodendrocyte | 52% | 61% |

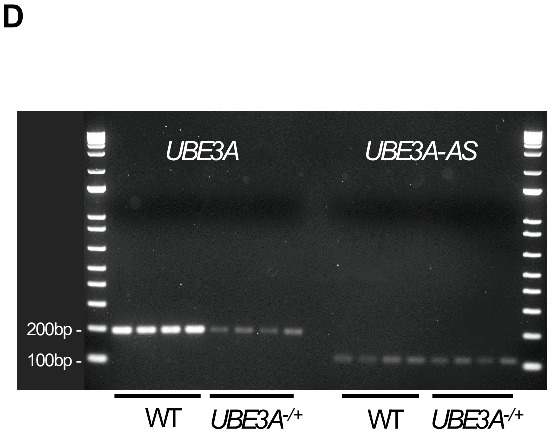


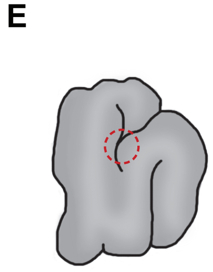


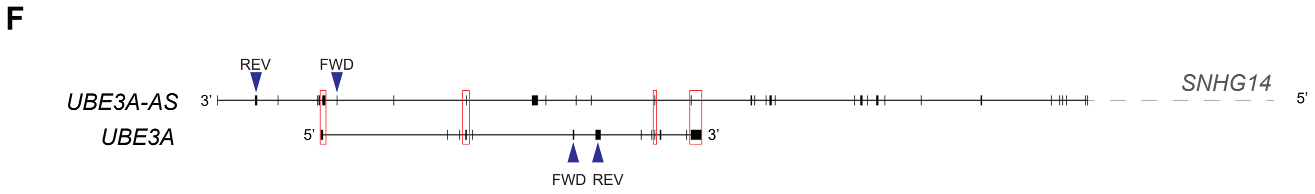


**Supplementary Figure 1. A custom annotation was necessary to detect the overlapping pig *UBE3A* and *UBE3A-AS* reads using the cellranger pipeline**

**A. UMAP of pig frontal cortex with original Sus scrofa 11.1 reference genome with cell type labels.** The original pig reference genome was generated using *Sus scrofa* 11.1 fasta and unaltered RefSeq GTF GCF_000003025.6 with cellranger mkref. Identical computational settings as described in the main methods were applied.

**B. WT versus *UBE3A*$^{-/+}$ heatmap expression of UBE3A in the analysis using the original pig reference.** Without annotation of *UBE3A-AS* and removal of the overlapping exons, multi-mapping reads are not accurately identified and are included in *UBE3A* read counts. Since *UBE3A-AS* expression is specific to the neuronal clusters, this inaccurately raises the *UBE3A* counts in neuronal clusters, making it appear that there is more *UBE3A* expression in the *UBE3A*$^{-/+}$ neurons. Per 10X Technical Note CG000376, 50% of intronic reads from brain nuclei are 5' ("antisense") in the 3'v3.1 chemistry preventing strandedness from being inferred in this context. Removing overlapping exons of the *UBE3A* and *UBE3A-AS* references and counting only exonic mapping reads removes multi-mapping reads, albeit at the cost of reducing overall *UBE3A* read depth.

**Supplementary Figure 1 (Continued). A custom annotation was necessary to detect the overlapping pig *UBE3A* and *UBE3A-AS* reads using the cellranger pipeline**

**C. Relative expression of *UBE3A* by cell type between the original pig reference versus the custom annotation with overlapping exons removed.**
*UBE3A* read fractions (*UBE3A*$^{-/+}$/WT) in non-neuronal cells are not dramatically altered, demonstrating that the incidental loss of *UBE3A* read-depth does not unduly bias *UBE3A* measurements. With the exclusion of multimapping reads, *UBE3A* expression is proportional to expected values.

**D. Agarose gel of *UBE3A* and *UBE3A-AS* PCR products.** PCR products run on an agarose gel (from same qRT-PCR primer sequences specified in Main Figure 1H) from frontal cortex tissue of the same eight animals used in the single nuclei RNA-sequencing indicate that the primers used are specific to their targets. *UBE3A* product length 193nt; *UBE3A-AS* product length 111nt.

**E. Diagram of the frontal cortex punch taken from the left hemisphere.** The brain was cut into 4mm coronal sections. The second 4mm section was used for taking a punch for each animal.

**F. Pig *UBE3A* and *UBE3A-AS* transcripts.** The pig *UBE3A-AS* transcript is not annotated in the current Ensembl or RefSeq gene annotations. Strand-specific bulk RNA-sequencing on poly-A enriched RNA isolated from the pig cortex was used to generate a consensus gene model of the *UBE3A-AS* transcript using methods as previously described [1]. Like the human and mouse *UBE3A-AS/Ube3a-AS* transcripts, we found that the pig *UBE3A-AS* transcript is transcribed across the paternal *UBE3A* allele, spliced, and 3'-polyadenylated. Overlapping *UBE3A* and *UBE3A-AS* exons (in red boxes) were removed from the custom annotation to ensure specificity of read counts. The GTF file of the custom *UBE3A-AS* annotation is included in the SRA accession. Blue arrows indicate primer sequences (Supplementary File 11) used in Main Figure 2E and Supplementary Figure 1F. The forward *UBE3A-AS* primer spans exons. Coordinates of the overlapping exons are: chr1:141984591-141987465; chr1:141974996-141975160; chr1:141925289-141925484; chr1:14188718-141887772.

Abbreviations: Uniform Manifold Approximation and Projection = UMAP; Wildtype = WT; *UBE3A* maternal deletion = *UBE3A*$^{-/+}$

#### Supplementary Figure 2

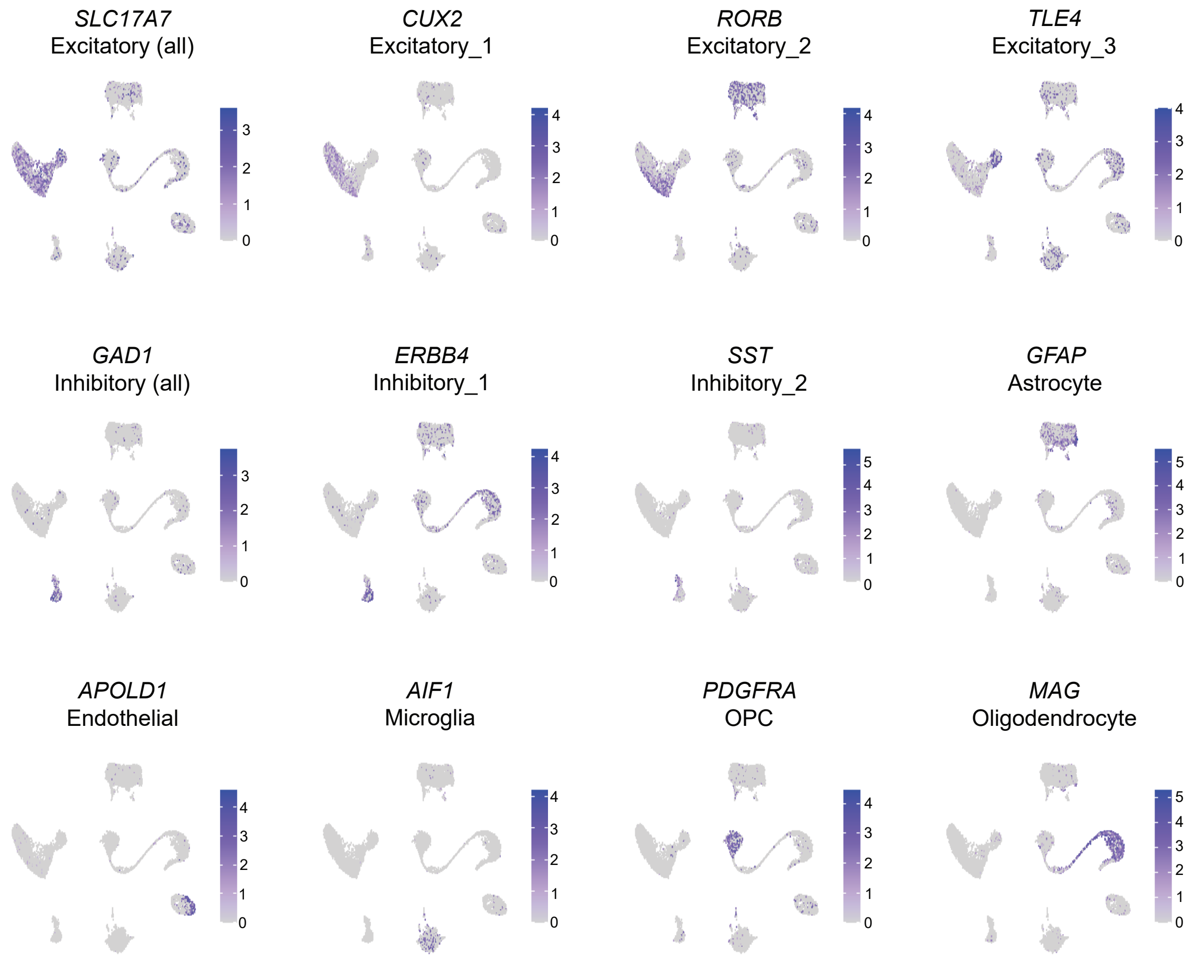


**Supplementary Figure 2. Heatmap expression of cell type marker genes.**

Representative marker genes are *SLC17A7* (pan Excitatory); *CUX2* (Excitatory_1); *RORB* (Excitatory_2); *TLE4* (Excitatory_3); *GAD1* (pan Inhibitory); *ERBB4* (Inhibitory_1) expression corresponds to parvalbumin+ inhibitory neurons cluster, however parvalbumin reads were low. *SST* (Inhibitory_2); *GFAP* (Astrocyte); *APOLD1* (Endothelial); *AIF1* (Microglia); *PDGFRA* (OPC); *MAG* (Oligodendrocyte).

**Supplementary Figure 3**

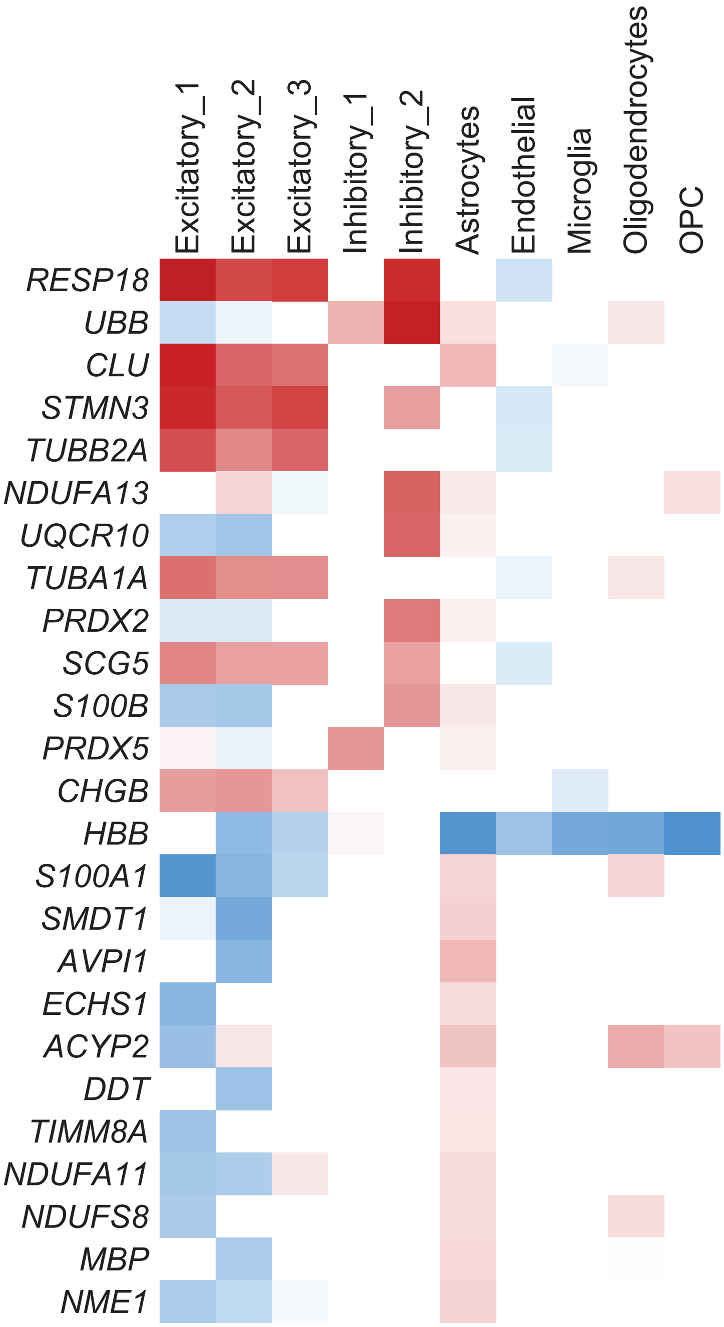


**Supplementary Figure 3. Genes with dual patterns of expression.**

The top 25 genes are displayed that are both significantly upregulated and downregulated in different cell types.

#### Supplementary Figure 4

A

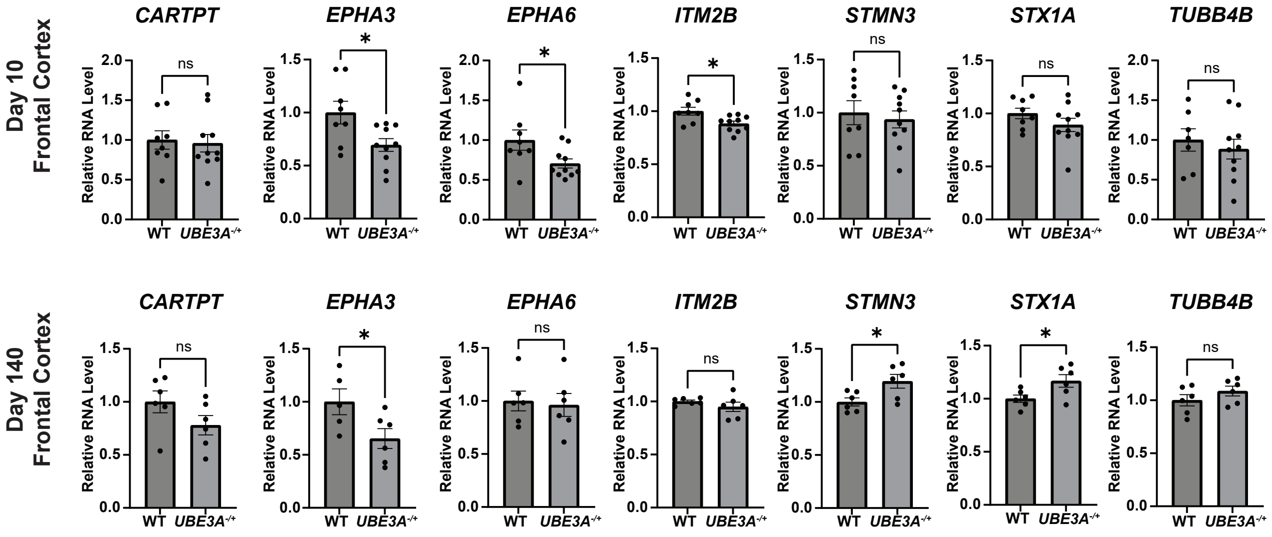


B

Bulk RNA-sequencing Volcano Plot

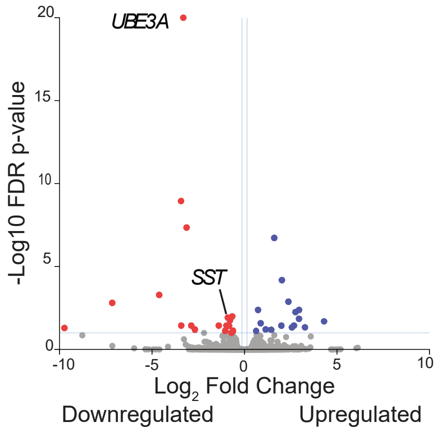


C

35 DEGs in Bulk RNA-Sequencing
FDR < 0.1

| Gene Name | Fold change | FDR p-value |
|---|---|---|
| ***UBE3A*** | -9.93 | 6.083E-74 |
| *LMO4* | -10.77 | 1.142E-09 |
| *LOC100127131* | -8.79 | 4.47E-08 |
| *KLHL40* | 3.05 | 1.904E-07 |
| *RHBDL2* | 4.07 | 0.000066 |
| *SDHAF4* | -24.42 | 0.000520 |
| *TH* | 5.20 | 0.00134 |
| *SLC5A12* | -142.39 | 0.00157 |
| ***CXCL2*** | 7.70 | 0.00412 |
| *MT-ND6* | 1.66 | 0.00412 |
| *GIPC3* | 6.77 | 0.00563 |
| ***PER1*** | -1.59 | 0.0101 |
| ***SST*** | -1.66 | 0.0120 |
| *ZBTB16* | -1.87 | 0.0120 |
| *WNT8B* | 7.68 | 0.0141 |
| *NLGN4X* | -1.73 | 0.0178 |
| *SCD* | 19.66 | 0.0202 |
| ***APOLD1*** | 1.83 | 0.0261 |
| ***GPNMB*** | -7.38 | 0.0361 |
| *EGR2* | -1.96 | 0.0361 |
| *ENSSSCG00000042700* | -2.62 | 0.0361 |
| *NAGLU* | -10.75 | 0.0361 |
| *ENSSSCG00000041294* | 6.36 | 0.0361 |
| *GJA9* | 3.99 | 0.0372 |
| *FOSB* | -1.79 | 0.0376 |
| *SMIM32* | 9.66 | 0.0461 |
| *TTR* | 5.91 | 0.0463 |
| *ENSSSCG00000040321* | -852.53 | 0.0515 |
| *ENSSSCG00000050388* | 2.21 | 0.0622 |
| *ENSSSCG00000032301* | -6.41 | 0.0622 |
| *IAPP* | 2.72 | 0.0661 |
| *NOS1* | -2.07 | 0.0749 |
| *ARC* | -1.54 | 0.0749 |
| ***WSB1*** | 1.54 | 0.0774 |
| *COL25A1* | -1.63 | 0.0990 |

D

RNA-sequencing DEGs Overlap

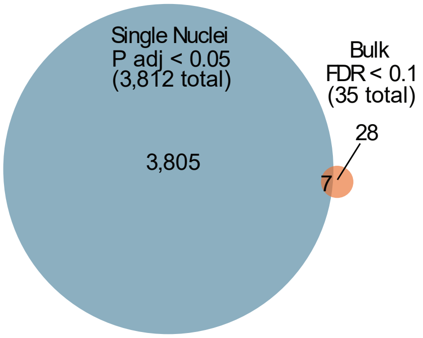

**Supplementary Figure 4. Orthogonal validation of differential expression results in qRT-PCRs and bulk RNA-sequencing.**

**A. qRT-PCR results for a subset of genes differentially expressed in the single cell RNA-sequencing.** Genes were examined in 10-day-old (top row) and 140-day old (bottom row) animals. For 10-day-olds, data includes the 8 animals used in the single nuclei RNA-sequencing and their littermates. Mean averages of these genes differ by less than 25%, revealing relatively small differences in expression. With the small changes in mean expression, the use of an assay which is not cell type-specific, and small sample size, it is difficult to validate the results observed in the single cell RNA-sequencing. *ITM2B* demonstrates a small reduction in expression at day 10, but this was not confirmed at day 140, likely due to higher variation and smaller sample size. *CARTPT*, *STMN3*, *STX1A* are not different in the day 10 results, but greater differences are observed in the day 140 animals, indicating greater disruptions may occur with age. *EPHA6* is reduced in the qRT-PCR data but was upregulated in the single nuclei RNA-sequencing data, showing an expression pattern in the opposite direction. TaqMan assays utilized: *RPL4* (ribosomal protein L4) was used as the reference gene: Ss03374067_g1; *CARTPT* (cocaine and amphetamine-regulated transcript protein): Ss03386009_u1; *EPHA3* (ephrin type-A receptor 3): Ss04321636_m1; *EPHA6* (ephrin type-A receptor 6): Ss06837582_m1; *ITM2B* (integral membrane protein 2B): Ss03380123_s1; *STMN3* (stathmin 3): Ss06898811_m1; *STX1A* (syntaxin 1A): Ss06936429_mH; *TUBB4B* (tubulin beta 4B class IVb): Ss06936831_g1.
**B. Volcano plot of bulk RNA-sequencing differential expression analysis.**
**C. Table of DEGs identified in bulk RNA-sequencing. Differential expression results on adolescent (140 day-old) pig frontal cortex** yielded 35 DEGs (FDR p-value < 0.1). Several genes (including *WNT8B*[2, 3], EGR2[4, 5], *IAPP*[6, 7], *NOS1*[8], *KLHL40*[9]) have previously been implicated in disorders with similarities to Angelman syndrome such as autism, Rett syndrome, Prader Willi syndrome, and myopathy. We also observe dysregulation of genes previously associated with Angelman syndrome including *ARC*[10] (activity-regulated cytoskeleton-associated protein) and *PER1*[11, 12] (period circadian regulator). Gene names bolded are also differentially expressed in the single nuclei analysis, including *SST* (somatostatin).
**D. Venn diagram illustrating overlap between bulk RNA-sequencing DEGs with the single nuclei DEGs**. A total of 3,812 DEGs (Bonferroni-adjusted P-value < 0.05) were revealed in the single nuclei RNA-sequencing data, of which only 7 overlapped with the 35 DEGs (FDR P-value < 0.1) of the bulk RNA-sequencing.

#### Supplementary Figure 5

**A**

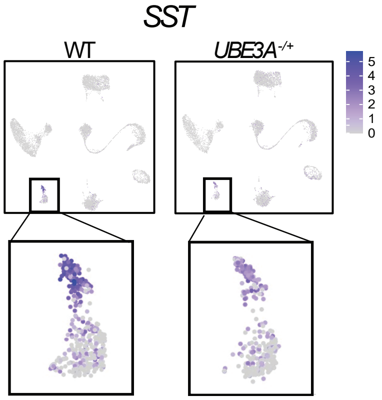


**B**

Somatostatin Receptors Differential Expression by Cluster

| | | avg_log2FC | p_val_adj |
|---|---|---|---|
| Excitatory_1 | *SSTR1* | -1.24 | 1.17E-12 |
| Excitatory_1 | *SSTR2* | -1.07 | 4.68E-09 |
| Excitatory_1 | *SSTR3* | 0.78 | 2.27E-06 |
| Excitatory_2 | *SSTR3* | 0.69 | 0.0079 |

**Supplementary Figure 5. Somatostatin and its receptors are differentially expressed in the neurons of the pig frontal cortex in single cell RNA-sequencing.**

**A.** UMAP of *SST* expression with zoomed panels of the inhibitory neurons.
**B.** Table of clusters where *SST* receptors were differentially expressed.
